# Urea amidolyase as the enzyme for urea utilization in algae: functional display in *Chlamydomonas reinhardtii* and evolution in algae

**DOI:** 10.1101/2024.05.16.594469

**Authors:** Honghao Liang, Senjie Lin, Yuanhao Chen, Jingtian Wang, Muhammad Aslam, Jing Chen, Hong Du, Tangcheng Li

## Abstract

Urea is a crucial nitrogen nutrient source for algae with the potential to stimulate harmful algal blooms, but the molecular machinery underpinning urea uptake and assimilation by algae is not fully understood. Urease (URE) is commonly regarded as the responsible enzyme, but the urea amidolyase (UAD) system, albeit known to exist, has hardly been studied. Here, the phylogenetic distribution, expression patterns, and functional roles of UAD system are examined, which comprises subunits *DUR1*, *DUR2*, and *DUR3*. We find a widespread occurrence of UAD, spanning four major phytoplankton lineages, and potentially independent evolution of URE and lineage-specific loss. Besides, a stronger regulation of UAD by environmental nitrogen concentrations compared to URE is uncovered in both global ocean and local dinoflagellate-dominant bloom events. CRISPR-based mutation in *Chlamydomonas reinhardtii* shows that subunit *DUR2* is essential for urea utilization. *DUR2* inactivation led to completely growth restriction and upregulation of *DUR1* and *DUR3A*, suggesting its functional interaction with them. In contrast, *DUR3B* inactivation only partially halted urea uptake and cell growth but significantly reduced gene expression across the entire UAD system. These findings not only reveal the crucial role of *DUR2* in urea utilization in *C. reinhardtii* and potentially in many other algae, but also suggest *DUR2* to be a more suitable indicator of urea utilization than urease, and underscore the importance to consider both URE and UAD enzyme systems when urea utilization by algae is assessed.

## Introduction

Phytoplankton play a pivotal role in the global carbon cycle, contributing to approximately 50% of global carbon dioxide fixation [1]. This process not only provides essential organic carbon to ecosystems but also generates oxygen, supporting the entire global biota. However, the growth and productivity of phytoplankton are critically dependent on availability of nitrogen (N), which is required for their metabolic processes [2]. In nature environments, N exists in various forms, with the most common being nitrate, nitrite and ammonium, serving as the primary sources for algae, but they only account for 2% to 11% of total dissolved nitrogen in most aquatic habitats [3–5]. Therefore, dissolved organic nitrogen (DON) is an important nitrogen source for algal growth, with urea contributing 50% or more of the total nitrogen used by planktonic communities [6–8]. Urea concentration can reach as high as 150 micromolar (μM), accounting for over 50% of the total dissolved N pool [9, 10]. With the development of human agricultural culture, furthermore, urea fertilizer use has increased 100-fold worldwide in the past four decades [11]. Excessive inputs of urea has led to eutrophication or nutrient regime shift thus influencing the composition of algal communities in aquatic ecosystems [9, 10]. Therefore, studying the ability and molecular mechanisms by which algae utilize urea is of significant importance for the management of eutrophic water bodies and the protection of aquatic ecosystems. The ability and characteristics of algae to utilize urea have been extensively studied, however, the underlying mechanisms, which potentially differ among lineages of algae [7, 12, 13], are poorly understood and underexplored.

Two enzymes systems, urease (URE) and urea amidolyase (UAD), exist in algae [14, 15]. Some species possesses one of these whereas others, such as fungi *Fusarium oxysporum*, cyanobacteria *Microcoleus vaginatus* FGP-2 contains both systems [16, 17]. URE is a nickel-dependent metalloprotein that is typically composed of three structural subunits (*ureA*, *ureB*, and *ureC*) and four accessory proteins (*ureD*, *ureE*, *ureF*, and *ureG*) [17, 18]. Among them, *ureC* was usually used as the target gene for URE analysis because it is the largest of the genes encoding urease functional subunits and contains highly conserved regions [19]. It is better characterized and is known to widely present in bacteria, fungi, plants, dinoflagellate, diatoms, and more [17, 20]. The measurement of URE activity can be used to estimate the potential contribution of urea to the nitrogen demand of phytoplankton and bacteria [21, 22]. In contrast, UAD is an ATP- and biotin-dependent protein [23]. It possesses two active subunits (urea carboxylase, *DUR1*; allophanate hydrolase, *DUR2*) and is only previously found in few fungal species, green algae, and bacteria [24, 25]. Hence, the significance of UAD in the process of urea utilization in aquatic environments is frequently underestimated when compared with URE [7, 17]. In addition, previous studies have shown that URE and URD enzyme systems do not coexist in the same species [26, 27], portraying a mutually exclusive evolutionary trend.

Regardless of what enzyme mediates urea degradation, phytoplankton capable of utilizing urea must use a passive or active mechanism to transport urea, including aquaporins or urea/amide channel proteins and *DUR3* protein [28]. *DUR3* is one of the high-affinity urea transport proteins, which includes three subunits *DUR3A*, *DUR3B* and *DUR3C* in the green alga *C. reinhardtii* but can be different in other algae. High-affinity nutrient transporters are commonly induced by nutrient deficiency. For example, the upregulated expression of *DUR3A*, *DUR3C* has been detected in *C. reinhardtii* under nitrogen deficiency conditions [29]. In addition, expression of *UAD* was significantly increased under nitrogen-deficient and urea-enriched environments, while the expression of *URE* was predominantly upregulated only under nitrogen deficiency [30, 31]. Several studies even indicated that URE genes are not regulated by changes in nitrogen availabilities (see http://cyanoexpress.sysbiolab.eu/) [17], suggesting that UAD and URE have different regulatory strategies in response to fluctuated nitrogen environments. Previous studies have shown that the induction of *DUR3* in *Candida albicans* may potentially require UAD [30]. Thus, we speculate that there could be an intricate relationship and coordinated expression between *DUR3* and urea-utilizing enzymes. However, still relatively little is known about their connection.

To address the gap of research regarding how the UAD and URE systems are distributed taxonomically, we searched the genomes of various species of representing all major phyla algae for the presence of URE and UAD systems. Next, we mined the gene catalogue derived from the *Tara* Oceans expedition (MATOU) to investigate the occurrence and expression of the URE (*ureC*) and UAD (*DUR1* and *DUR2*) genes across phytoplankton species in the global ocean [32]. We further examined the evolutionary trajectory of the two urea-utilizing mechanism by conducting phylogenetic analyses. Moreover, to prove the role of UAD in urea utilization, we focused on *C. reinhardtii* and knocked out *DUR2* and *DUR3B* using CRISPR/Cas9, which has been successfully used in studying *C. reinhardtii* as well as diatoms [33–36]. *C. reinhardtii* is commonly used as a model organism for studying gene functions because of a well-characterized genome, simple cell structure, and fast growth rate [37, 38]. Our results will reveal the significance of UAD in the urea cycle of aquatic environments, and shed light on how phytoplankton adapt to urea-enriched aquatic environments through the UAD system.

## Materials and methods

### Microalgal strains and culture conditions

*Chlamydomonas reinhardtii* wild strain 137c (cc-125) and *DUR3B* mutant (LMJ.RY0402.103895) were obtained from the *Chlamydomonas* Resource Center (https://www.chlamycollection.org/). The *DUR2* mutants were generated in this study using CRISPR gene knockout. All *C. reinhardtii* cultures were maintained in modified TAP medium at a temperature of 25℃ under a square-wave 14 h: 10 h light: dark regime, with a light intensity of 80 μmol m^-2^ s^-1^. The TAP medium was modified by combining NH_4_ ^+^ and urea as nitrogen sources in different molar ratios to achieve a total nitrogen concentration of 7 mM. The liquid medium was continuously shaken at 110 rpm using an orbital shaker (SK-O180-Pro, Dragon Lab, China). Experimental cultures were set up in Erlenmeyer flasks, each containing 0.15 L of TAP medium with three nitrogen conditions: N deficient condition (denoted as N-), 7 mM NH_4_ ^+^ (denoted as A), and 3.5 mM urea (denoted as U). The cell concentration was determined every two days throughout the experiment using a hemocytometer (YA0811, Solarbio, China). The specific growth rate (μ) was calculated using formula of μ=(lnX_1_-lnX_0_)/(t_1_-t_0_), where X_1_ and X_0_ represent the cell concentrations at times t_1_ and t_0_, respectively. To determine the urea utilization capacity of wild-type (WT) and mutant (*m*DUR2 and *m*DUR3B) strains, each strain was diluted to 5×10^5^ cell mL^-1^, 5×10^4^ cell mL^-1^, and 5×10^3^ cell mL^-1^ and cultured on 1.5% (*w/v*) agar plates which contained either NH_4_ ^+^ or urea or NH_4_ ^+^ + paromomycin, respectively.

### sgRNA design and cleavage activity verification in vitro

The single guide RNAs (sgRNAs) targeting the *DUR2* gene were designed by Benchling (benchling.com), resulting in two high on-target score sgRNA sequences (*DUR2*-1: 5’-CAAGTAGCTTGTCCACTGCG-3’ and *DUR2*-2: 5’-CTACCACCAGAGATGAACCG-3’) in the first and second exon of *DUR2*. The sgRNAs were prepared in vitro as described by Yu et al. (2017). The oligos used for sgRNAs synthesis were summarized in Table S1. To verify the efficiency of designed sgRNAs, the *DUR2* target loci was amplified from genomic DNAs and then purified using the Universal DNA purification kit (DP214, Tiangen Biotech Co., China). For formation of the CRISPR ribonucleoprotein (RNP), the Cas9 protein (200 ng) purified from *E. coli* rosetta containing the plasmid (Addgene ID: 47327) and sgRNA (600 ng) were mixed together, and then incubated at 37℃ for 15 minutes. For target DNA cleavage reactions, 200 ng target DNAs were added to CRISPR RNP with a final volume of 20 μL and then incubated at 37℃ for 1 hour. Cleavage reactions were purified by the DNA purification kit (DP214, Tiangen Biotech Co., China) and separated by 1.5% agarose gel electrophoresis.

### *C. reinhardtii* transformation and mutations selection

Approximately 2×10^7^ cells were harvested and suspended in 1 mL of Max Efficiency Transformation Reagent (Thermo Fisher Scientific, USA) which containing 40 mM sucrose. Prior to electroporation, cells were shaken at 300 rpm under 40℃ conditions. SpCas9 (100 μg, 0.53 nmol) and sgRNA (60 μg, 1.6 nmol) were mixed at 37℃ for 15 min to assemble CRISPR RNP complexes. The exogenous double stranded (donor) DNA, which contained 45 bp *DUR2* specific homology arms and paromomycin cassette (including RBCS2 promotor, an aph*VIII* reporter gene and RBCS2 terminator) that were amplified from pCrU6.4-SaCas9cloning/aph*VIII* (Chlamydomonas Resource Center ID: pPH339) plasmid. Then, about 5 × 10^6^ cells, RNP complexes and 2 μg of donor DNA were added to 4 mm gap cuvette (Bio-rad) and incubated on ice for 5 min. The cells were then transformed via an electric pulse (600 V, 50 μF, infinity resistance) on a Bio-Rad Gene Pulser Xcell Electroporator (Bio-Rad) [39]. Immediately after electroporation, the cells were suspended in 10 mL TAP medium containing 40 mM sucrose for recovery. Finally, the cells were cultured in low light conditions (20 μmol m^-2^ s^-1^) at 25 °C for 24 h, after plated on TAP agar plate containing 10 μg mL^-1^ of paromomycin.

After 8 days of incubation, single clones were picked and transferred to 96-well plates containing 180 μL TAP medium and 10 μg mL^-1^ paromomycin per well. After all wells turned green (about 4 days), 40 μL of cells were taken from each well to extract genomic DNA using the Phire Plant Direct PCR Master Mix (Thermo Fisher Scientific, USA). Then, colony PCR was conducted to verify the target site under the thermal cycles of 98℃ for 5 min, followed by 35 cycles of 98℃ for 5 s, 55℃ for 30 s, and 72℃ for 30 s. The PCR products were separated by electrophoresis on a 1% agarose gel, and the target bands were sequenced to confirm the indel mutations. All the primers sequences used in this study were summarized in Table S1.

### Western blot analysis of *DUR2* protein accumulation

Cells of WT and *DUR2* mutant strains, cultured under urea as the only nitrogen supply, were harvested by centrifugation at 8000 × *g* for 10 min and then resuspended in RIPA Lysis Buffer (Beyotime Institute of Biotechnology, China). The protein concentration was determined using the BCA Protein Assay Kit (TransGen Biotech, Beijing, China). Approximately 20 μg of protein were separated on a 10% (*w/v*) SDS-PAGE gel and subsequently transferred onto Immobilon P membrane (EMD Millipore, Billerica, MA). Immunoblots were performed to detect the presence of *DUR2* using a custom-made antiserum specific to *DUR2* (GenScript Biotech Corp., Nanjing, China) at a dilution of 1:2000. As a reference, the abundance of the β-subunit of ATP synthase (Atpβ) was assessed using a polyclonal antibody (AS05-085, Agrisera AB) at a dilution of 1:3000. The selection of the Atpβ protein as a reference is grounded in its previously reported stable abundance in *C. reinhardtii* [40]. After three washes in TBS buffer for 10 min each, the membranes were incubated with a secondary antibody (mouse anti-rabbit IgG-HRP, Santa Cruz Biotechnology) diluted 1:3000 for 1 hour. Next, the membranes were washed and treated with BeyoECL Star Kit (Beyotime Institute of Biotechnology, China) to detect the immunoreactive bands on the Amersham Imager 600 system (GE Healthcare, Boston, MA).

### Pigment contents, *F*v/*F*m ratio, dry weight, total cellular carbohydrate contents and extracellular urea concentration

The pigment contents of C. *reinhardtii* in each treatment group were determined spectrophotometrically. Briefly, 5 × 10^6^ cells were harvested and washed three times with ddH_2_O. The cell pellets were then resuspended in 5 mL of methanol and incubated at 4℃ overnight in the dark. After centrifugation at 5000 × *g* for 5 min, the supernatant was used to determine the absorption spectra using a UV2400 spectrophotometer (SOPTOP, China). The calculation of Chl *a* and Chl *b* was performed according to the published equations [41]. To determine the *F*v/*F*m ratio of each culture, culture samples were taken and incubated in darkness for 30 min and measured on the Imaging-PAM (91090, Heinz Walz GmbH, Germany).

Approximately 1 × 10^8^ cells were filtered onto Whatman GF/C glass filters (Whatman International Ltd, Maidstone, UK) and washed with ddH_2_O. Dry weight was determined using gravimetric analysis, as described by Vasileva et al. (2020). Total cellular carbohydrate content was determined using the anthrone-sulfuric acid method Sánchez Mirón et al. (2002). The concentration of remaining urea in the media was measured photometrically using diacetylmonoxime method [44] on a UV2400 spectrophotometer, (SOPTOP, China) at wavelength of 520 nm.

### RNA isolation and sequencing

Previous studies have shown that a complete set of genes including *DUR1*, *DUR2*, and *DUR3A* reach their maximum expression at 2 to 4 h after transfer to N-free medium [29]. Therefore, samples were collected at 4 h after transferring cells into different nitrogen treatment conditions in this study. The cells were harvested by centrifugation at 2,500 × *g* at 4℃, following a previously reported protocol [45]. Total RNA was isolated using TRIzol reagent (MRC, Cincinnati, USA) coupled with Direct-zol™ RNA MiniPrep kit (Zymo Research, Irvine, CA, USA). Only samples with the RNA Integrity Number (RIN) ≥ 7 were used for RNA-seq libraries construction. Each library was sequenced on an Illumina HiSeq xten/NovaSeq 6000 at Majorbio Bio-pharm Technology Co., Ltd (Shanghai, China). The reads were then mapped to the *C. reinhardtii* genome (Creinhardtii_v5_6) using HISAT2 [46]. The RSEM and DESeq2 software were used to estimate transcript abundances and identify differentially expressed genes (DEGs) between WT and mutant strains [47, 48]. KEGG enrichment analysis was performed with KOBAS 2.1.1 [49].

### Evolutionary analyses and biogeographic distribution of selected genes

To investigate the evolutionary trend of *DUR1*, *DUR2* and *ureC* genes among representative bacteria, fungi and algae, a species phylogenetic tree was constructed based on TIMETREE database (https://timetree.org/). The abundance of coding gene of *DUR1*, *DUR2* and *ureC* was downloaded from NCBI Database and JGI PhycoCosm database. To explore the prevalent and species taxonomy of *DUR1*, *DUR2* and *ureC* in the global scale, the *DUR1* domain (Pfam ID: PF02682), *ureC* domain (Pfam ID: PF00449), and the full length of *DUR2* gene of *C. reinhardtii* were used as queries to retrieve these genes from the Ocean Gene Atlas (OGA) database [32], following previously reported methods [50]. The reason for using the full-length of the *DUR2* gene is that its domain is an amidase, and the presence of amidase in many enzymes would lead to large number of false sequence alignment results. Then, the eukaryote-enriched Marine Atlas of the *Tara* Ocean Unigenes (MATOU) with the cut off e-value of 1E^-3^ were queried in OGA, resulting in an output including alignment results, homolog sequences FASTA files, and normalized abundance of the homologs. Subsequently, NR database and eggNOG-mapper search were performed to re-annotate the retrieved homologs. Only hits annotated as *DUR1*, *DUR2*, or *ureC* coupled with the e-value < 1E^-8^ were selected for subsequent analyses [50]. Gene abundances were computed as RPKM (reads per kilobase covered per million of mapped reads), and the distribution pie chart was calculated by using ‘percentage of total abundance per gene’. The biogeographic distribution was plotted in R (v. 4.1.1) using ggplot2 packages. Furthermore, environmental data was also obtained in the Ocean Gene Atlas (OGA) database (Table S7 and S8), and mantel test was performed by the OmicStudio tools (https://www.omicstudio.cn/tool) to analyze the correlation between the gene expression and environmental factors.

### Statistical analysis

To assess the statistical significance of differences between the WT and mutant strains in different treatment groups, the SPSS software (version 20.0, IBM, US) was used in this study. Before comparison of different treatment groups, all data were first tested for homoscedasticity and normality. One-way analysis of variance (ANOVA) was performed, followed by post-hoc comparisons using Tukey’s honestly significant difference tests. For comparisons involving cell number and medium urea concentrations, a generalized linear model repeated measure procedure was conducted. This procedure tests both the growth parameter and treatment-time factors, employing a two-way analysis of variance (ANOVA). All data were presented as mean ± standard deviations with three biological replicates (N = 3), and statistical significance was determined at a significance level of p < 0.05 (* or different letters above the columns).

## Results

### Phylogenetic tree and geographic distribution for UAD and URE systems

To understand how originally both UAD and URE occurs in phytoplankton, the phylogenetic tree was constructed based on TIMETREE database (Fig. 1A). The results showed that the ancestral cyanobacteria *Gloeobacter violaceus* contained UAD but not URE system, suggesting the UAD may have originated earlier than URE in phytoplankton. Furthermore, UAD and URE seem to have undergone independent evolution as they showed unlinked presence and absence in different taxa. Besides, the domains of key subunit *DUR1* and *DUR2* were very conserved among major phyla algae but showed lineage-specific loss (Fig. 1B, C). The species distribution results showed that the *DUR1* and *DUR2* were predominantly distributed in Cyanobacteria, Fungi, Rhodophyta, Chlorophyta and Dinophyta, whereas URE (*ureC*) was mainly distributed in ‘red type’ algal lineages such as Cyanobacteria, Fungi, Rhodophyta, Chlorophyta, Dinophyta, Haptophyta, Bacillariophyta, Cryptophyta, Eustigmatophyta, Phaeophyta, Prasinodermohyta, indicating a broader distribution of the URE system than UAD system in phytoplankton (Fig. 1A; Fig. S1). In addition, we found one cyanobacterial species, one fungus and one dinoflagellate contain both UAD and URE systems.

**Fig 1.**
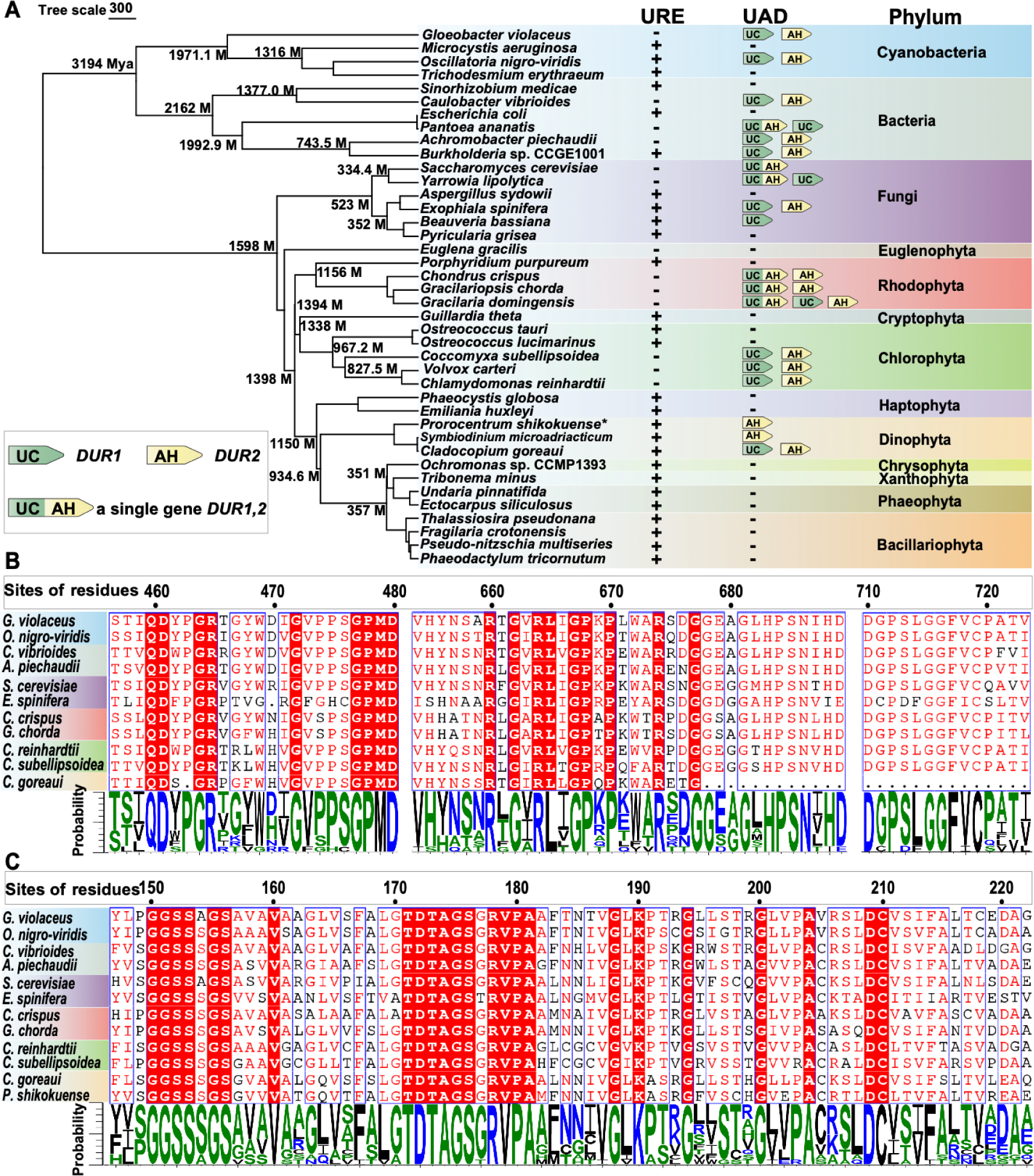
Evolutionary dynamics of DUR gene cluster and sequence conservation of DUR1 and DUR2. **(A) Urea carboxylase (*DUR1*), allophanate hydrolase (*DUR2*) and urease (*ureC*) among selected bacteria and algae mapped to the phylogenetic tree. (B) Alignment of conserved domain in DUR1 protein. (C) Alignment of conserved domain in DUR2 protein.** The phylogenetic tree was constructed based on TIMETREE database (https://timetree.org/). Lineage divergence time is indicated at each branch point; M, million years ago; asterisk indicates that the data were predicted by transcriptome. Probability was figured by https://weblogo.threeplusone.com.

The analysis of the *Tara* Ocean data yields 572, 258 and 258 identified hits of *DUR1*, *DUR2* and *ureC* respectively from most of the sampling stations, suggesting their overall prevalence in marine phytoplankton (Fig. 2A-B; Tables S3-S6). Besides, *ureC* showed a broader taxonomic distribution than *DUR1* and *DUR2* as the former were found in nine classified lineages, one unclassified lineage, and one cluster of unknown eukaryotes whereas DUR1 and DUR2 were found in six classified lineages, one unclassified lineage and one cluster of unknown eukaryotes (Fig. 2B). The transcriptomic expression of *DUR1*, *DUR2*, and *ureC* differed largely depending on the water depth and sampling station (Fig. 2A). Briefly, the more extensively distributed and relative higher abundance of *DUR1*, *DUR2*, and *ureC* were found on surface (SRF) than in the deep chlorophyll maximum (DCM) depth of the ocean. In addition, the abundance of *DUR1* in SRF and DCM seems to be higher than that of *ureC*, accounting for 50.27% and 53.90% of the total abundance respectively, while the *DUR2* was lower than *ureC*, accounting for only 20.83% *and* 16.73% (Table S3). On a single cell scale, the transcripts of *DUR1* (TPM = 14.74) were higher than *DUR2* (TPM = 4.37) in *C. reinhardtii* (WT) under urea as sole nitrogen environments in our transcriptomic data (Table 1), potentially suggesting that the UAD system is probably regulated by the expression of *DUR2*.

**Fig 2.**
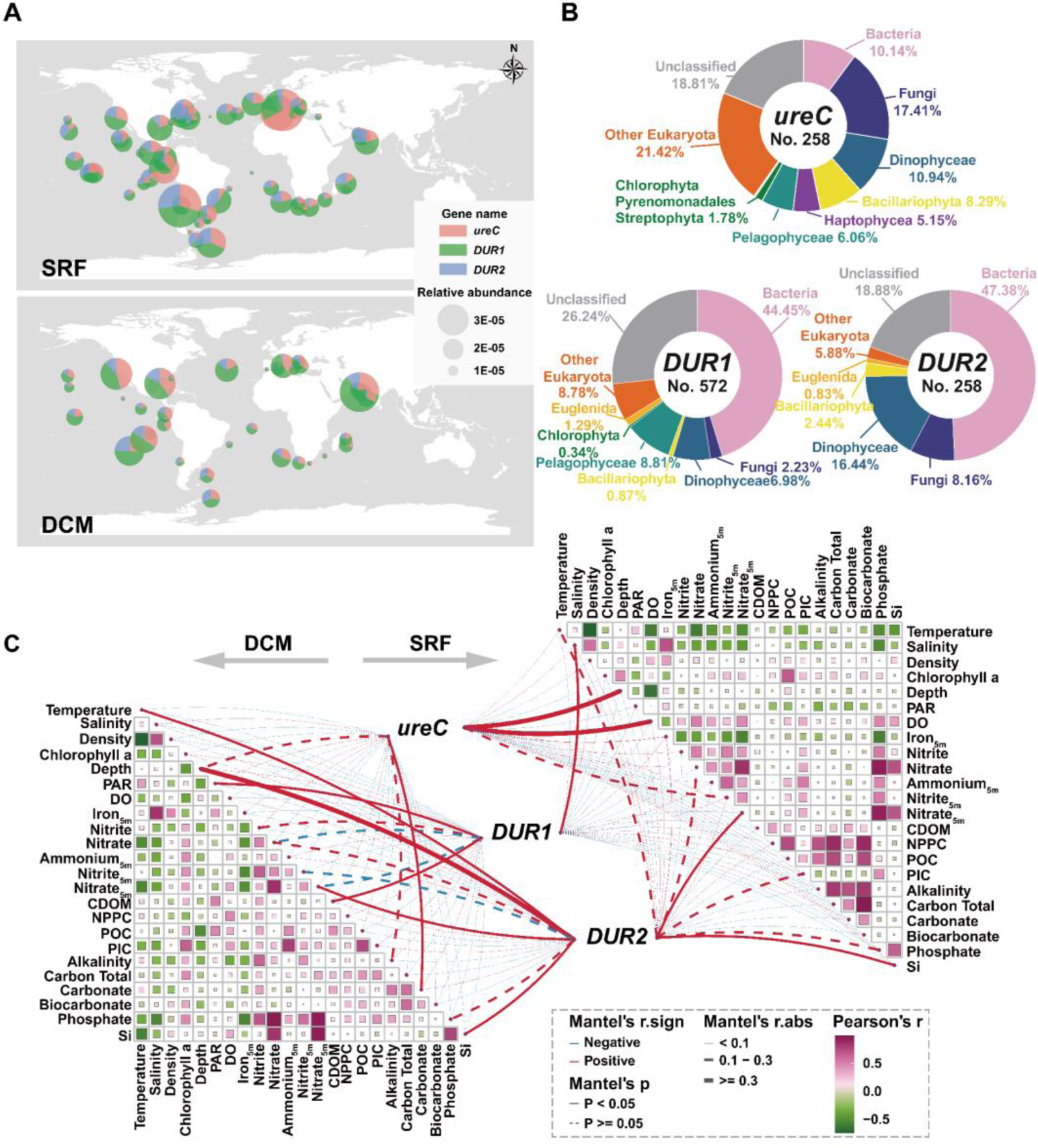
Biogeographic distribution, species taxonomy and interactions with environmental factors of *DUR1*, *DUR2* and *ureC* in *Tara* Ocean database. **(A)** Geographic distribution of *DUR1*, *DUR2* and *ureC* genes in the global ocean. SRF: layer of surface; DCM: the deep chlorophyll maximum. All data used in this analysis are listed in Table S3 to S6. **(B)** species taxonomy of *DUR1*, *DUR2* and *ureC* genes from (A); **(C)** Interactions across the expression of *DUR2*, *DUR1* and *ureC* with environmental factors. The thickness of the line represents the correlation between environmental factors and gene expression, and the color of the lines represents the significance level of the correlation. The intensity of the fill color of the box represents the correlation of the selected environmental factor from green (negative interaction), white, to purple (positive interaction). PAR: radiation, photosynthetically active per day; DO: dissolved oxygen; CDOM: colored dissolved organic matter; NPPC: net primary production of carbon; POC: particulate organic carbon; PIC: particulate inorganic carbon.

**Table 1.** The mean TPM value of urea transporters (*DUR3A*, *DUR3B* and *DUR3C*) and urea amidolyase (*DUR1* and *DUR2*) in WT, *m*DUR2 and *m*DUR3B under three nitrogen treatment conditions.

| Gene | WT_A<sup>*</sup> | WT_U<sup>#</sup> | WT_N-<sup>\$</sup> | *mDUR2*_U | *mDUR2*_N- | *mDUR3B*_U | *mDUR3B*_N- |
| --- | --- | --- | --- | --- | --- | --- | --- |
| *DUR3A* | 0.00 | 16.29 | 0.00 | 279.64 | 1.38 | 0.00 | 0.00 |
| *DUR3B* | 5.82 | 7.10 | 10.06 | 17.41 | 12.58 | NA | NA |
| *DUR3C* | 0.04 | 4.10 | 0.05 | 11.74 | 0.07 | 0.02 | 0.01 |
| *DUR1* | 0.14 | 14.74 | 5.32 | 199.50 | 12.09 | 2.71 | 4.00 |
| *DUR2* | 0.36 | 4.37 | 2.79 | NA | NA | 1.72 | 2.22 |

Note: “NA” indicates the TPM not available; “\*” represent ammonium condition, “#” represents urea condition and “\$” represents the nitrogen deficient conditions.

### Correlation analysis of UAD and URE with environmental factors across global ocean

Our analysis also revealed significant correlations between the expression of three genes and various environmental parameters in both the global and local ocean (Fig. 2C; Fig. S2). Notably, among the three genes, *DUR2* exhibited a more pronounced relationship with nitrate concentration (5 m water depth, 0.1 < r < 0.3; P < 0.05) compared to *ureC* (r < 0.1; P > 0.05) in both SRF and DCM layers across global ocean, suggesting a potential regulatory role of nitrogen nutrients on UAD system. Additionally, *DUR2* showed a significant relationship with silicon (Si) in the SRF layer, as well as temperature, depth, and Si in the DCM layer (P < 0.05). Although *DUR1* showed a strong relationship with nitrate and nitrite (0.1 < r < 0.3), the P-value was over 0.05. Moreover, *ureC* exhibited weaker relationships with environmental parameters, excluding depth and dissolved oxygen (DO) in the SRF layer, and carbonate levels in the DCM layer (Fig. 2C).

In the local scale, we analyzed the relationship of UAD and URE systems with environmental nitrogen on the shelf (C6 station) and slope (C9 station) of the South China Sea and Tongxin Bay of the East China Sea [51, 52]. Same as findings in the *Tara* Oceans database, *DUR2* exhibited a significant relationship with nitrate concentration (P < 0.05) but urease did not (P > 0.05) in both sampling sites, indicating a stronger regulation of the UAD system by environmental nitrogen concentrations compared to the URE system (Fig. S2).

### Knockout of *DUR2* gene in *C. reinhardtii*

To better understand the functional importance of the *DUR2*, we knocked out the *DUR2* gene in *C. reinhardtii*. Our CRISPR/Cas9 genome editing protocol was first shown to be effective through an in vitro experiment (Fig 3A, B), which indicated that one of the two sgRNAs (*DUR2*-1) was more efficient than the other (*DUR2*-2). *DUR2*-1 was used in the subsequent in vivo knockout experiment (Fig. 3B). After transformation, 12 mutant strains were obtained (12/68, 17% success rate), and among them 7 strains with a single band over 2200 bp from PCR amplification of *DUR2* were selected for Sanger sequencing (Fig. 3C; Fig S3). The sequencing results confirmed successful insertional mutation of the *DUR2* in two strains that prevented expression of the gene (Fig. 3D; Table S1). Furthermore, western blot analysis of these two mutants (*m*DUR2-21 and *m*DUR2-44) did not detect clear bands corresponding to *DUR2*, confirming the abolishment of *DUR2* gene translation (Fig. 3E). Therefore, *m*DUR2-21 and *m*DUR2-44 strains were used to verify the functional loss of *DUR2* and resultant phenotypes in *C. reinhardtii*.

**Fig 3.**
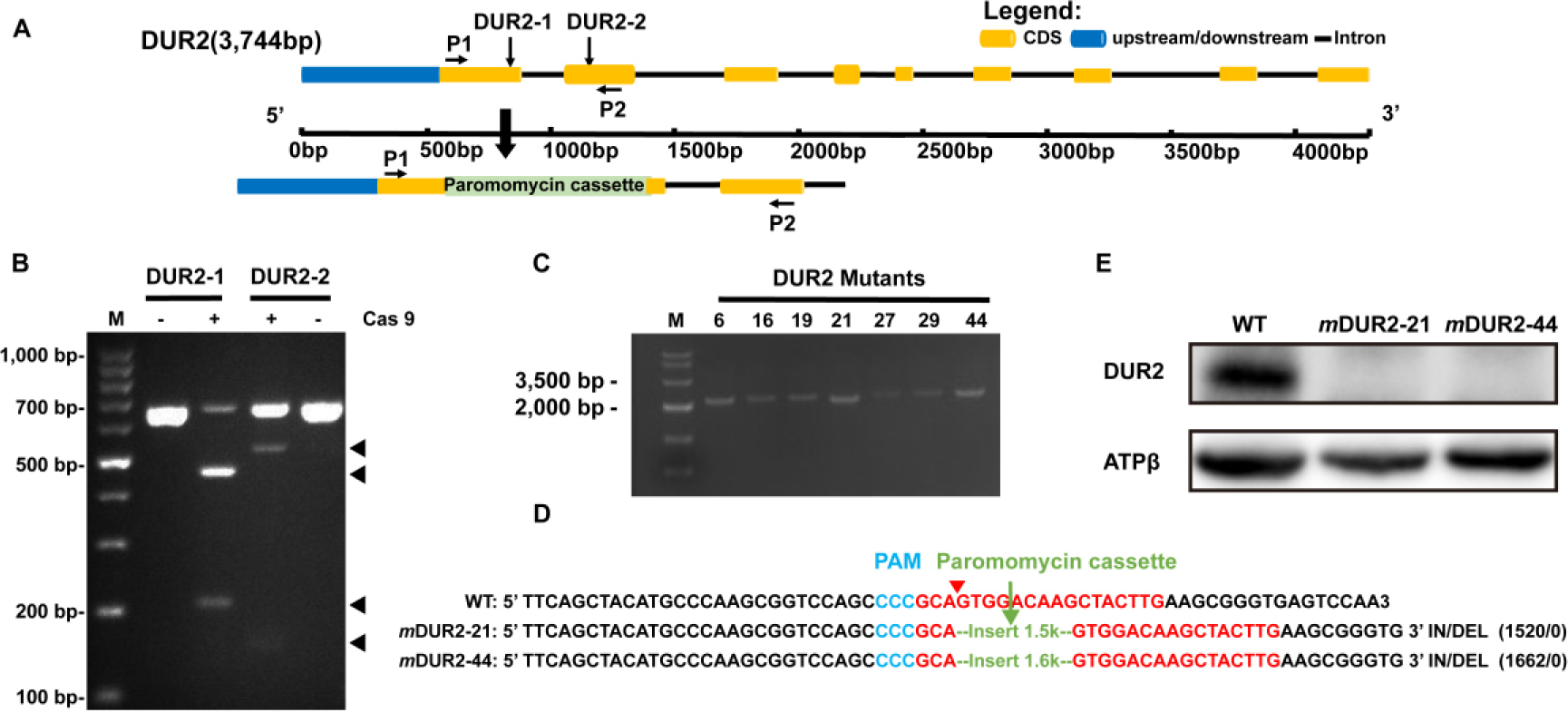
Successful knockout of *DUR2* gene in *Chlamydomonas reinhardtii*. **(A)** Design of sgRNA sequences for the *DUR2* gene. Locations of two sgRNAs for DUR2-1 and DUR2-2 are indicated by vertical thin arrows. P1 and P2 delineate the locations of the primers designed for amplifying the DUR2 target region. **(B)** Target region of *DUR2* gene amplified with specific primers, with the black arrowhead showing a deletion mutant band. M: DNA marker. **(C)** PCR analysis of seven selected clone after transforming CRISPR RNP. **(D)** Sequence alignment of representative mutant clones with the wild type (WT). The paromomycin cassette sequences are labeled in green; PAM sequences are labeled in blue; the sgRNA sequences are labeled in red. **(E)** Immunoblot image of *DUR2* protein and beta subunit of ATP synthase.

### The growths of *m*DUR2, *m*DUR3B and WT under three nitrogen environments

The growths of *m*DUR2-21, *m*DUR2-44, *m*DUR3B, and WT strains showed that the two *m*DUR2 strains and *m*DUR3B grew well under NH_4_ ^+^ amended with paromomycin, indicating that knockout of *DUR2* and *DUR3B* did not negatively impact growth of *C. reinhardtii* when NH_4_ ^+^ was supplied as the source of N nutrient (Fig. 4). In contrast, the growth patterns and cell colors of *m*DUR2, *m*DUR3B and WT were clearly different when urea was used as sole N source (Fig. 4A-B). The growth of *m*DUR2-21 and *m*DUR2-44 was strongly suppressed when urea was sole N source (0.012 ± 0.022 compared to 0.157 ± 0.018 in WT, P < 0.05), while that of *m*DUR3B was less suppressed albeit significantly slower than NH_4_ ^+^ condition (0.487 ± 0.022 versus 0.607 ± 0.007, P < 0.05; Fig. S4). In accordance, *DUR2* mutation completely abolished the ability to take up urea whereas *DUR3B* mutation only reduced urea uptake, as shown by the different temporal trends of urea concentration in the medium (Fig. 4C; P < 0.05, two-way ANOVA). This resulted in a negative correlation between the cell concentration and extracellular urea concentration for the wild type (R^2^ = 0.92, P < 0.05) but not for *m*DUR2 (Fig. 4D). These results indicate that the functional loss of *DUR2* and *DUR3B* genes deprived *C. reinhardtii* of the ability to grow on urea as the N source.

**Fig 4.**
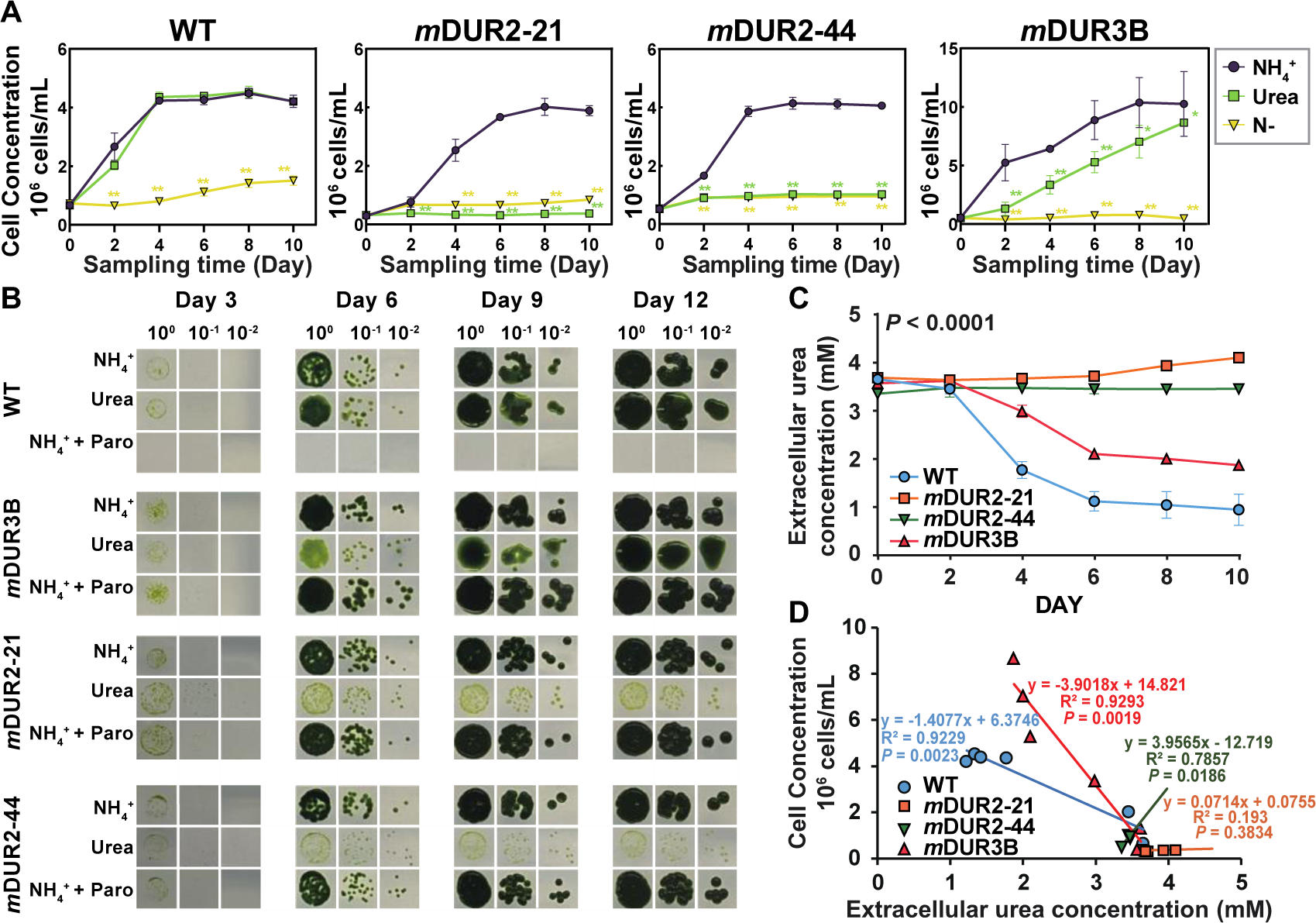
Change of growth and urea consumption after DUR2 and DUR3B mutation. (A, B) The growth of *C. reinhardtii DUR2* and *DUR3B* mutants grown under different nitrogen conditions on liquid and TAP mediums. (C) The concentration of urea left in the culture medium. (D) The relationship between cell concentration and extracellular urea concentration. Paro represents paromomycin; 10^0^, 10^-1^ and 10^-2^ represent cell concentration in 5×10^5^ cell mL^-1^, 5×10^4^ cell mL^-1^, and 5×10^3^ cell mL^-1^.

### The chlorophyll, *F*v/*F*m ratio, dry weight and carbohydrate of *m*DUR2, *m*DUR3B and WT

In general, between growing on NH_4_ ^+^ and urea, WT grew similarly well and showed no differences in chlorophyll, dry weight, carbohydrate content and *F*v/*F*m ratio (P > 0.05, Fig. S5). For the two *m*DUR2 strains, cellular chlorophyll content was 38.05% and 25.09% lower when growing on urea than when growing on NH_4_ ^+^ (P < 0.05, Fig. S5A). Although *m*DUR3B can grow under urea as only nitrogen condition, a significant decrease in chlorophyll was also observed in the urea culture group than NH_4_ ^+^ culture groups (P < 0.05). However, the *F*v/*F*m ratio showed no decrease in *m*DUR2 and *m*DUR3 growing on urea (Fig. S5B). Interesting, the maximum dry weight of the two *m*DUR2 strains were both observed in the urea-grown culture group (587.71 μg/10^6^ cells in *m*DUR2-21 and 252.25 μg/10^6^ cells in *m*DUR2-44), while the minimum was found in the NH_4_ ^+^-grown culture group (108.83 μg/10^6^ cells and 116.96 μg/10^6^ cells) (Fig. S5C). Similarly, mirroring the dry weight trend, the NH_4_ ^+^ group exhibited the lowest carbohydrate content, despite having the highest growth rate, suggesting that cell growth was enhanced at the cost of carbohydrate content (Fig. 4A; Fig. S5D).

### Shift of overall gene expression profiles in *m*DUR2 and *m*DUR3B

RNA-seq yielded 63,488,905 clean reads on average from each of the 27 cultures (Table S2). Approximately 91.67% of the clean reads were uniquely mapped to the reference genome and covered 93.66-96.80% of genomic predicted proteome (Table S2). Using a fold change ≥ 2 and an adjusted P-value < 0.05 as the criteria, 946 difference expression genes (DEGs) and only one significant KEGG pathway (nitrogen metabolism) were obtained from *m*DUR2_A vs. WT_A comparison groups, indicating *DUR2* gene knockout did not seriously influence the growth and gene regulation under NH_4_ ^+^ conditions (Fig. S6). In contrast, when grown on urea, a total of 8,313 DEGs were identified between the strains, 2115, 6628, 2080 and 869 DEGs obtained from *m*DUR2_U vs. WT_U, *m*DUR3B_U vs. WT_U, WT_N- vs. WT_U and WT_A vs. WT_U comparisons, respectively (Fig. 5A, B). Furthermore, only 111 DEGs were shared by the four comparison groups, revealing largely unique regulatory strategies after *DUR2* and *DUR3B* gene knockout (Fig. 5B). For *m*DUR2, 2,115 DEGs were significantly enriched in thirty KEGG pathways including ribosome, photosynthesis, glycolysis, porphyrin, chlorophyll metabolism, nitrogen metabolism, purine metabolism, one carbon pool by folate, pentose phosphate pathway, pyruvate metabolism, glycerolipid metabolism, biotin metabolism and carbon fixation, indicating widely intracellular regulation by *DUR2* (P < 0.05; Fig. S7).

**Fig 5.**
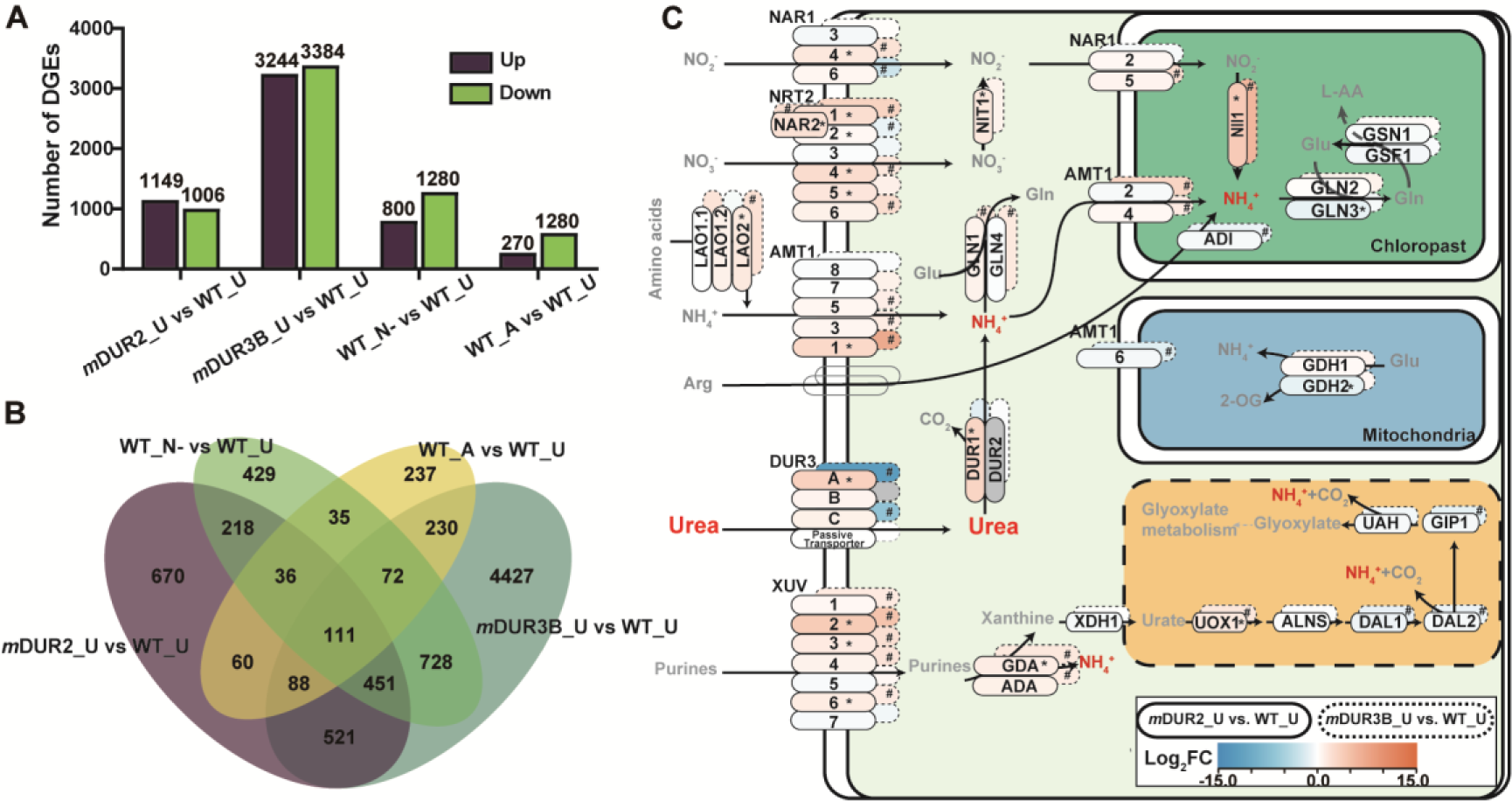
Shift of transcriptomic profile resulting from DUR2 and DUR3B mutation. (A) Differential expression genes. (B) Venn diagram showing shared genes among four comparison groups. (C) Remarkable nitrogen metabolic reconfiguration due to *m*DUR2 and *m*DUR3B inactivation. WT_A, WT_U, and WT_N-represent wild type *C. reinhardtii* cultured in NH ^+^, urea, and nitrogen deficiency conditions; *m*DUR2_U and *m*DUR2_N-represent *DUR2* mutant strain cultured in urea conditions and nitrogen deficient conditions; *m*DUR3B_U and *m*DUR3B_N-represent *DUR3B* mutant strain cultured in urea and nitrogen deficient conditions. Asterisk and pound signs indicate the significant regulatory genes; red boxes indicate increased expression, blue color indicates decreased expression; grey color indicates knocked out genes; transparent boxes with gray contours indicate unidentified genes in this study.

### Regulation of genes involved in urea metabolism in *m*DUR2 and *m*DUR3B

Three high-affinity urea transporters (*DUR3A*, *DUR3B*, and *DUR3C*), one *DUR1* and one *DUR2* encoding genes were identified in the genome of *C. reinhardtii* [37]. Among them, *DUR3A*, *DUR3C*, *DUR1* and *DUR2* are hardly expressed (∼ TPM = 0) in the WT grown on NH_4_ ^+^, while all of them showed up-regulation after the cultures were transferred from NH_4_ ^+^ to urea environment (Table 1; Fig. 5C). Furthermore, *DUR1* and *DUR2* genes were not only regulated by nitrogen forms, but also by nitrogen concentration, as shown in their significant upregulation under nitrogen deficient conditions (Table S9), consistent with the stronger relationship between expression of *DUR1*, *DUR2* and environmental nitrogen concentration in *Tara* Ocean database (Fig. 2C). After *DUR2* inactivation, the transcripts of *DUR3A* and *DUR1* were significantly increased, from a TPM of 16.29 to 279.64 and 14.74 to 199.50, respectively when the *mDUR2* cultures were grown on urea as sole nitrogen source (Table 1). These translate to 11.23- and 8.45-fold increases, respectively, indicating these two genes may be partially inhibited by *DUR2* in the WT. Comparatively, *DUR3B* and *DUR3C* were markedly less influenced by *DUR2*, as their expression was only moderately up regulated, with a fold change between 1.5 and 2 (P < 0.05) in *DUR2* mutation (Table S9).

*DUR1* and *DUR2* transcripts peaked at ∼2 to 4 h and *DUR3B* at 12 to 24 h, exhibiting varied expression patterns in response to nitrogen fluctuations in *C. reinhardtii* [29]. In this study, the transcript abundance of *DUR3B* was higher than that of other four urea-related genes (*DUR3A*, *DUR3C*, *DUR1* and *DUR2*) under NH_4_ ^+^ environment in WT, and also showed relative stable after the cultures were transferred to urea (TPM from 5.8 to 7.1) or N-deficient conditions (TPM from 5.8 to 10.1), potentially indicating the *DUR3B* is less influenced by nitrogen chemical form or nitrogen concentration (Table 1). After *DUR3B* inactivation, *DUR3A*, *DUR3C* and *DUR1* gene transcripts were all reduced, which was consistent with the decreased urea absorption capacity of *m*DUR3B under urea as only nitrogen source conditions (Fig. 4C; Fig. 5C; Fig. S8). Collectively, the UAD system in *C. reinhardtii* is notably influenced by the chemical form of the environmental nitrogen source. Specifically, the inactivation of the *DUR2* promotes the expression of urea pathway-related genes, particularly *DUR1* and *DUR3A*, while the inactivation of *DUR3B* significantly inhibits the expression of the entire set of urea pathway-related genes (Fig. S9).

## DISCUSSION

Nitrogen is an essential nutrient for phytoplankton as it is a key component of amino acids, chlorophylls, and nucleobases biosynthesis [53, 54]. It is well recognized that phytoplankton species have evolved mechanisms to scavenge diverse source of nitrogen such as DON to cope with nitrogen deficient environments [55, 56]. Urea is an important DON component of aquatic ecosystems [7], and increasingly shown to promote harmful algal blooms [57]. URE is commonly regarded as the responsible enzyme, but the UAD system, albeit known to exist, has hardly been studied. Our functional genetic manipulation, transcriptome profiling, and physiological observations collectively revealed the critical role and some of the regulatory mechanisms of UAD system in *C. reinhardtii*. Furthermore, our analyses shed light on the distinct evolutionary trajectories of UAD and URE systems in algae. By mining the *TARA* Ocean meta-transcriptomic data, we found compelling evidence that both UAD and URE system are at play across the global ocean but being distributed among diverse phytoplankton lineages (Fig. 1, 2). The methodology employed in our study and the mutants generated serve as a fundamental foundation for enhancing our understanding of UAD and URE system and how algae employ UAD to adapt to fluctuations in aquatic environments.

### Evolution of algal UAD and URE systems and their relationship to environmental nitrogen

Prior to the discovery of UAD in bacteria, it was thought that the urea pathway exists only in some yeasts and green algae [7, 15]. In the present study, we found the UAD pathway in four major lineages of phytoplankton, including Chlorophyta, Dinophyta, Cyanobacteria, and Rhodophyta, and is distributed across a broad geographic range, which is more prevalent than generally appreciated (Fig. 1, 2). In addition, UAD exists in an ancient photosynthetic ancestor (*Gloeobacter violaceus*), which does not have URE, potentially suggesting a more ancestral evolutionary position of UAD than URE in phytoplankton. However, UAD seemed to be lost quickly in cyanobacteria and descendant algal lineages and gave way to URE, which dominated some of the ‘red type’ algal lineages such as diatoms, dinoflagellates, and phaeophytes (Fig. 1A). This is consistent with previous study that found genera *Gloeobacter* and *Gloeomargarita* in cyanobacteria that only contained UAD but likely diverged early from other cyanobacteria which contained the URE [17]. URE occurs in 12 higher plankton lineages, including Dinophyta, Cyanobacteria, Fungi, Chlorophyta and Rhodophyta, and some species in these five lineages contain UAD or both two urea utilization pathways (Fig 1A). For instance, we found one species contains both pathways such as a symbiodiniacean *Cladocopium goreaui* in Dinophyta. This observation challenges previous findings that urease and urea amidolyase activities were not present simultaneously in the same unicellular algae, being mutually exclusive in algae [26, 27].

The measurement of URE activity has been used to estimate the potential contribution of urea to the nitrogen demand of phytoplankton and bacteria [21, 22]. However, based on the *Tara* Oceans data and local meta-transcriptomic data, we found *urease* exhibited a less pronounced relationship with environmental nitrogen across the global ocean (Fig. 2C; Fig S2). Indeed, several studies performed with different cyanobacteria showed that the URE genes are not regulated by changes in nitrogen availabilities [17, 58]. However, UAD genes including *DUR1*, *DUR2*, *DUR3A* and *DUR3C* were significant regulated by the chemical form and concentration of nitrogen in *C. reinhardtii* and potentially in many other algae in the global ocean (Fig. 2C; Fig 5C; Fig S2). For instance, *DUR1* and *DUR2* showed significant relationships with environmental nitrogen in dinoflagellate-dominated algal blooms (Fig. S2C). These previous and current data suggest that UAD genes, especially *DUR2*, are more suitable indicators of urea utilization than URE gene. What’s more, the more abundant *DUR1* transcripts than *ureC* from phytoplankton identified in global ocean meta-transcriptomes potentially indicate a key position of UAD in the marine nitrogen cycle and underscore the importance to consider both URE and UAD enzyme systems when urea utilization by algae is assessed (Fig. 6).

**Fig 6.**
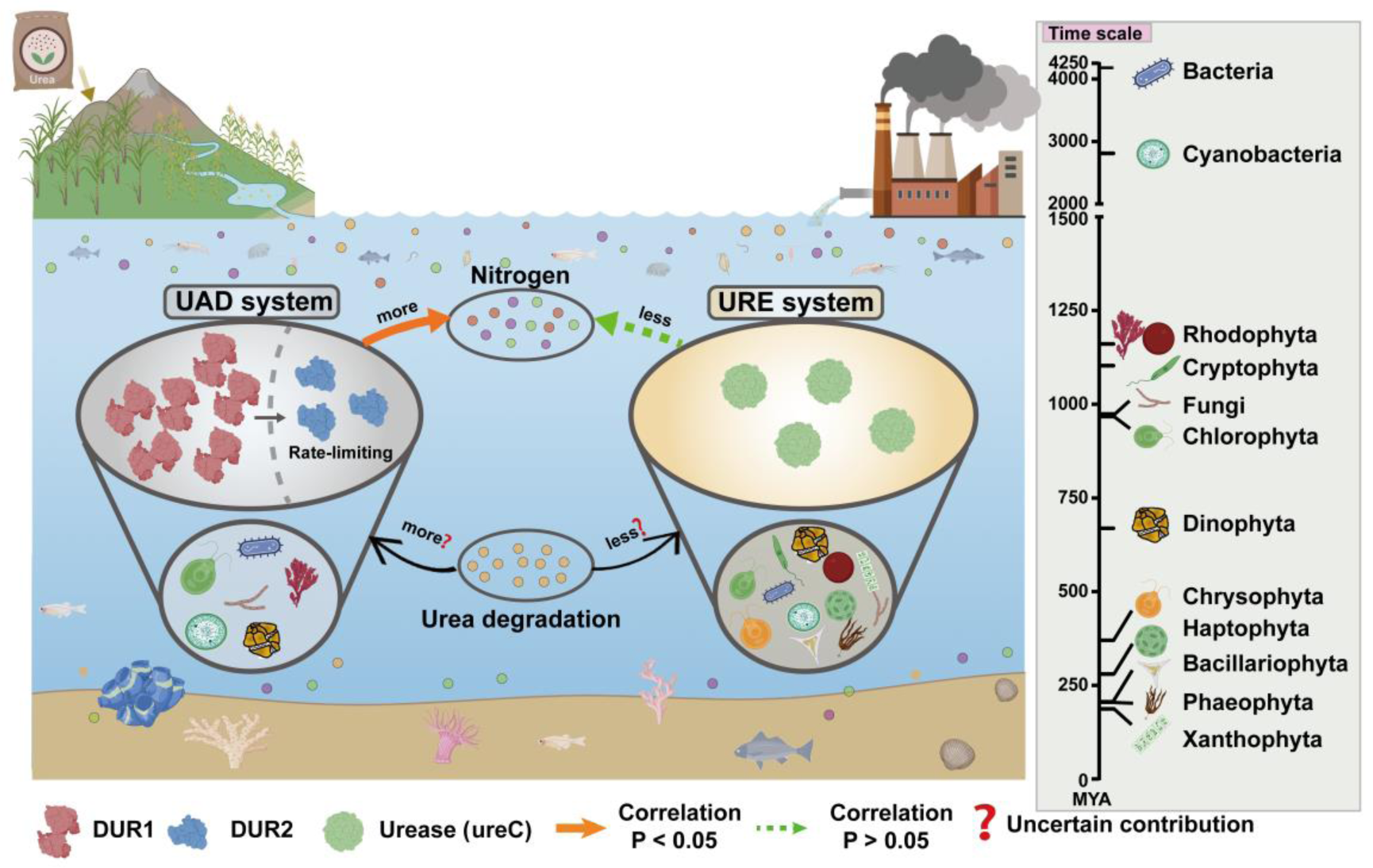
Schematic illustration of phylum distribution and its relationship to environmental nitrogen in UAD and URE systems. Different numbers of elements in UAD and URE systems represent the level of gene expression. Time scale was constructed based on TIMETREE database.

### Incongruent evolution of UAD subunits and potential functional coordination

The enzyme UAD was first characterized in the yeast *Candida utilis* and occur either as a fusion of urea carboxylase (*DUR1*) and allophanate hydrolase (*DUR2*), or as two separate proteins that work together to degrade urea, suggesting that, they may be closely related [7, 59]. However, previous studies reported that there is no substrate channeling or interdomain/intersubunit communication between *DUR1* and *DUR2*, and the catalytic efficiency of the two is independent of each other [60]. Interestingly, in some algal species, *DUR1* and *DUR2* occur as two domains on one protein (such as *Yarrowia lipolytica*) but in others they are present as separate genes (such as *Gracilaria domingenesis*) (Fig. 1A). Likely, there has been evolutionary events fusing the two functionally related domains into one gene. Gene fusion occurs frequently in eukaryotes such as the RAN binding protein 2 (*RANBP2*) gene in human [61]. The prevailing consensus suggests that gene fusion plays a pivotal role in intermediate transfer facilitation, enhancement of catalytic efficiency, and regulatory processes [62]. However, *DUR1* and *DUR2* function as independent entities seems prefer to the “selfish operon” model, clustering of functionally related genes serves simply to facilitate the successful inheritance of the gene cluster, both by horizontal gene transfer and by vertical transmission [60]. Notably, the gene fusion event observed in UAD is unique to Rhodophyta within the algal phylum, while in other algal species, UAD has evolved as an independent gene and seems lost at random (Fig. 1A) This observation suggests that the two subunits comprising UAD exhibit tendencies toward functional and evolutionary independence within algae.

On a global scale across a latitudinal gradient, *DUR1* was found to be more highly expressed than *DUR2* (Fig. 2). At the single-cell scale, we found that the transcripts of *DUR1* (TPM = 14.74) were 3.4 times higher than *DUR2* transcripts (TPM = 4.37) in *C. reinhardtii* under urea condition, potentially indicating that the *DUR2* may be an inducer or rate-limiting enzyme in UAD pathway. Indeed, we found that the cell completely lost the ability to utilize urea after *DUR2* inactivation under urea condition (Fig. 4). In addition, a previous study found that cell can hold a relative slower growth rate after urease inactivation in bacteria *Halomonas meridiana* [63], potentially suggesting the UAD pathway plays an irreplaceable role in urea utilization process.

Generally, *DUR2* is primarily associated with the decomposition of urea, specifically breaking down allophanate into NH_3_ and CO_2_ [30, 59]. In the present study, we found the *DUR2* can also regulate the urea transport of *C. reinhardtii* (Fig. 4B). After *DUR2* gene knockout, *C. reinhardtii* stop absorbing urea and the urea concentration is maintained at a stable concentration in the culture medium that might expand the knowledge of this common reaction in biological systems (Fig. 4B). Furthermore, we found *DUR2* inactivation resulted in *DUR3A* and *DUR1* expression upregulation. However, despite *DUR3A* upregulation, no urea transport into cells was detected in *m*DUR2. This conflicting result suggest potential existence of a feedback system between urea degradation and uptake. In addition, up-regulated the expression of *AMT* and *NRT* encoding genes were found in most phytoplankton under nitrogen deficient condition, e.g. *Fugacium kawagutii* [56], *Karenia brevis* [64] and *Prorocentrum shikokuense* [65]. Most genes encoding dissolved inorganic nitrogen transporter (e.g. *NRT* and *AMT*) and purine transporter (e.g. *XUV*) were upregulated in *m*DUR2 under urea as sole nitrogen environments (Fig. 6; Fig S9), suggesting a broad and coordinated molecular regulatory feedback on the deficiency of nitrogen nutrient as a result of urea utilization machinery breakdown.

### Differential responses of *m*DUR2 and *m*DUR3B to urea environment

Physiological performances and gene expression patterns revealed two distinct response strategies of *DUR2* and *DUR3B* under urea as sole nitrogen conditions (Fig. 4 to 5). The *DUR2* gene knockout resulted in complete inhibition of urea uptake and cell growth rate, while the *DUR3B* gene knockout only slowed down urea uptake and cell growth rate, indicating that the function of *DUR3B* can be complemented by a functional analog (Fig. 4B). *DUR3* is plasma membrane-localized high-affinity transporter composed of *DUR3A*, *DUR3B* and *DUR3C*, among which *DUR3A* and *DUR3B* are encoded on linkage group VIII while the *DUR3C* is encoded on linkage group XIX in *C. reinhardtii.* [66]. Therefore, the reason why the inactivation of the *DUR3B* gene does not completely inhibit urea absorption in cells may be due to its independent functioning from *DUR3A* and *DUR3C* such that when *DUR3B* is disrupted, *DUR3A* and *DUR3C* can partially fulfill the function. In addition, aquaporins such as *AtTIP1.1*, *AtTIP1.2*, *AtTIP2.1* and *AtTIP4.1* in *Arabidopsis thliana* and *NtTIPa*, *NtAQP1* in *Nicotiana tabacum* have been reported to participate in a low-affinity urea transport system [67–69]. Consistent with the possibility that aquaporins complement the function of *DUR3*, the genes for aquaporins (*MIP2* and *MIP1*) were indeed upregulated after *DUR3B* inactivation in *C. reinhardtii* (Table S10).

### Concluding remarks

This study serves as primer for employing CRISPR RNP genome editing technology to investigate the regulatory mechanisms of urea utilization through UAD in green algae *C. reinhardtii* and evolution in algae. The results indicates that the UAD system is more widely distributed than previous appreciated and may have originated earlier than URE system in phytoplankton. However, UAD seems to be lost quickly in cyanobacteria and descendant algal lineages and gave way to URE, which dominated some of the ‘red type’ algal lineages such as diatom and dinoflagellates. Besides, the UAD system shows a stronger correlation with environmental nitrogen form and nitrogen concentration than the URE system in both local and global surveys, potentially suggesting that UAD is a more suitable indicator of urea utilization than urease. Although the expression of *DUR1* subunit in the UAD system is higher in the global plankton transcriptome data, *DUR2* appears be an inducer or rate-limiting enzyme in UAD pathways of urea breakdown, as confirmed by near zero cell growth rate and cell absorption of urea after *DUR2* inactivation in *C. reinhardtii*. Notably, *DUR2* inactivation can upregulate the expression of *DUR1* and *DUR3A*, as evidenced by a significant increase in gene transcription following *DUR2* inactivation, suggesting the operation of a sophisticated feedback system in the urea utilization pathway. Collectively, this study unravels a crucial role of UAD system in phytoplankton urea utilization and highlights the *DUR2* in UAD system in *C. reinhardtii* and potentially in many other algae and delves into its ecological implications in adaptive mechanisms.

## Author contributions

Conceptualization: Tangcheng Li, Hong Du, Senjie Lin; Data curation: Honghao Liang; Formal analysis: Tangcheng Li, Honghao Liang; Funding acquisition: Tangcheng Li, Hong Du; Methodology: Tangcheng Li, Honghao Liang, Jingtian Wang, Yuanhao Chen, Jing Chen; Writing-original draft: Honghao Liang, Tangcheng Li; Writing - review & editing: Senjie Lin, Tangcheng Li

## Acknowledgement

We wish to thank Jianqing Lin, Yan Gao and Chongming Zhong from Shantou university for technical advice and assistance. This work was supported by the Natural Science Foundation of China grants (NSFC 42206116), the Guangdong Basic and Applied Basic Research Foundation (No. 2021A1515110001), Science and Technology Plan Projects of Guangdong Province (No. 2021B1212050025) and the Program for University Innovation Team of Guangdong Province (2022KCXTD008).

## Data availability statement

Raw sequencing data has been made available at NCBI in the SRA (Short Read Archive) database under accession number PRJNA1056916.

