## supplemental figures for "Urea amidolyase as the enzyme for urea utilization in algae: functional display in *Chlamydomonas reinhardtii* and evolution in algae"

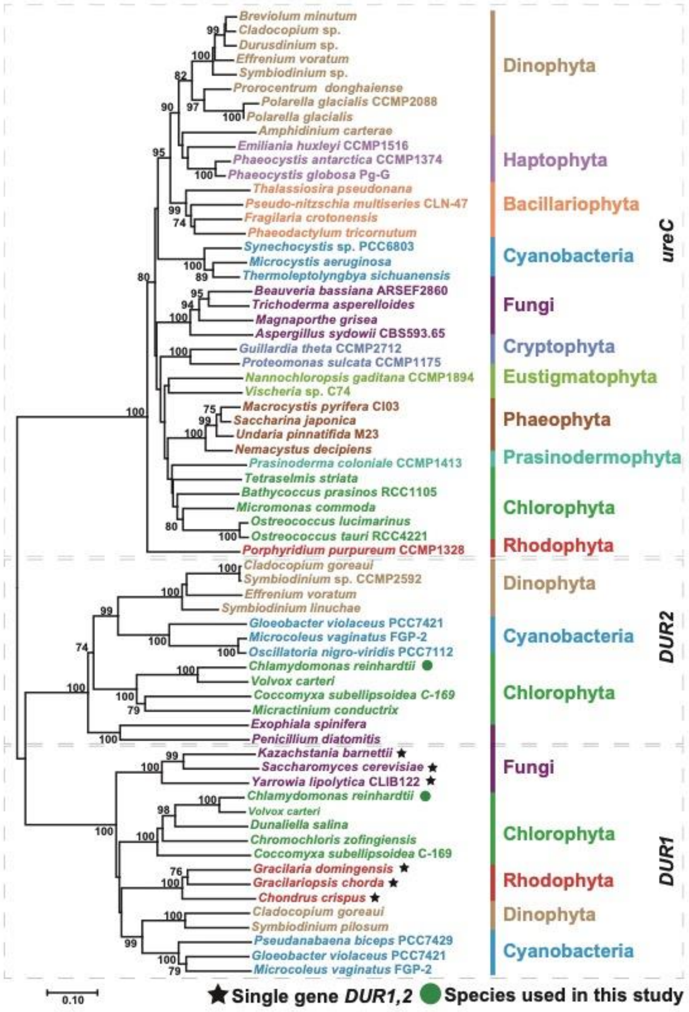


Figure S1. **Phylogenetic tree inferred from amino acid sequences of *DUR1*, *DUR2* and urease.** Only bootstrap values equal to or higher than 70% are shown. The pentagram represents the urea amidolyase sequence of the species was encoded by a single gene *DUR1,2*. The green dots represent the species used in this study.

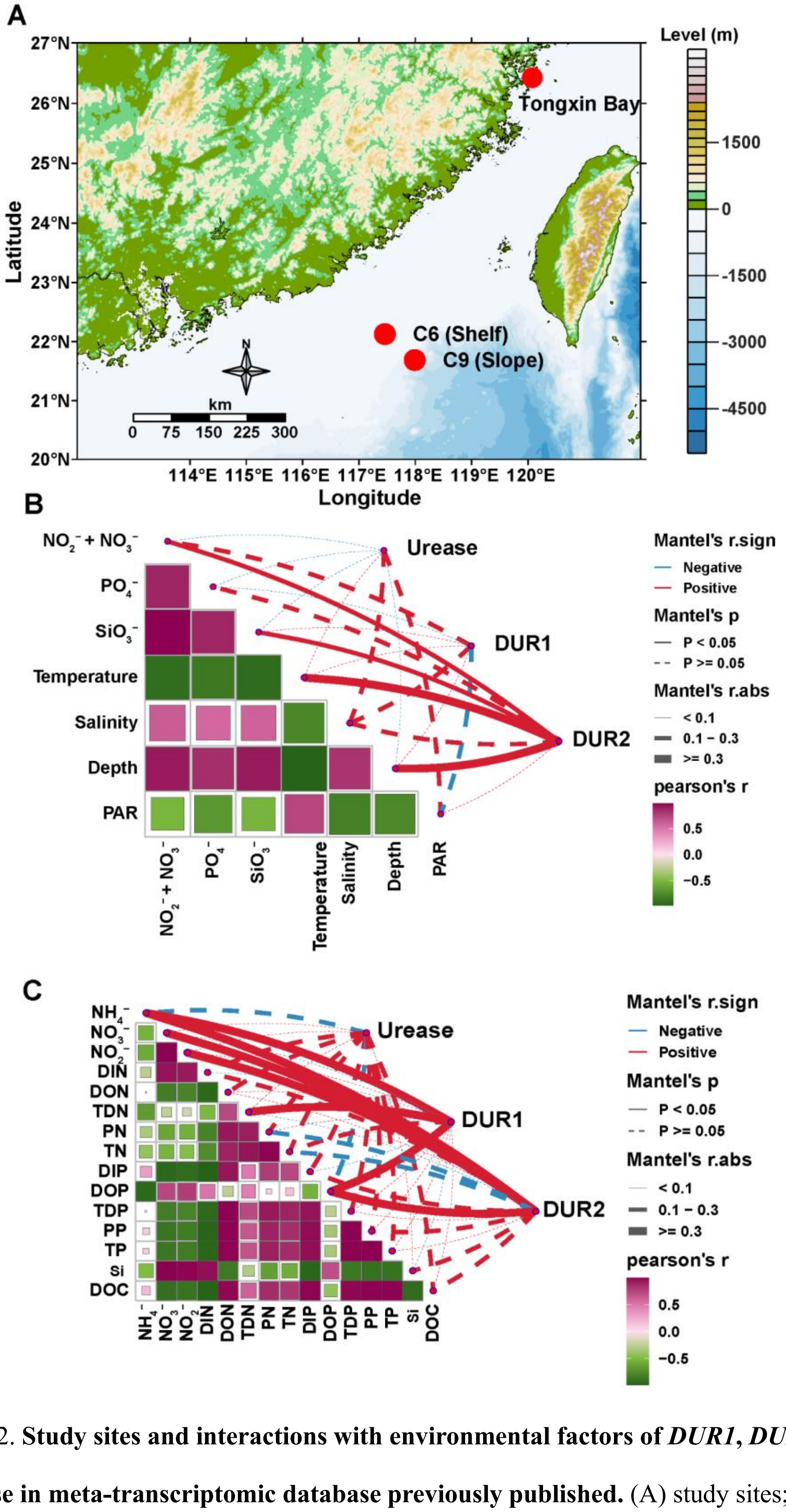


Figure S2. **Study sites and interactions with environmental factors of *DUR1*, *DUR2* and urease in meta-transcriptomic database previously published.** (A) study sites; (B) environmental samples collected from C6 and C9 stations [52]; (C) dinoflagellate-dominant algal bloom samples collected from Tongxin Bay of Fujian, China [51].

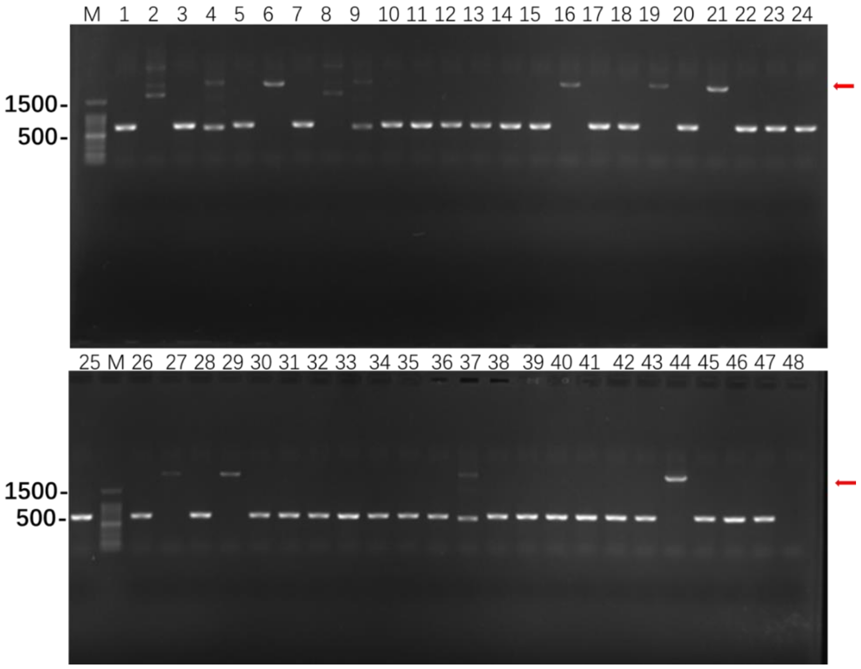


Fig. S3. PCR analysis of selected clones after transform CRISPR RNP. Lane 2.4.8.9 and 37 have more bands while the 6.16.19.21.27.44 have single bands. M: DNA marker.

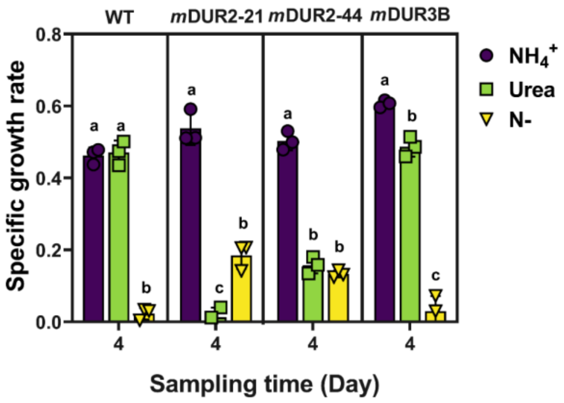


Fig. S4. The specific growth rate of four strains in different nitrogen medium at day 4. Significant differences of each strain in different treatment are indicated by different letters, ANOVA p <0.05 (a, b, c and d). Each strain cultured in $NH_4^+$ was used as control. Data are reported as the mean ± standard deviation (n = 3).

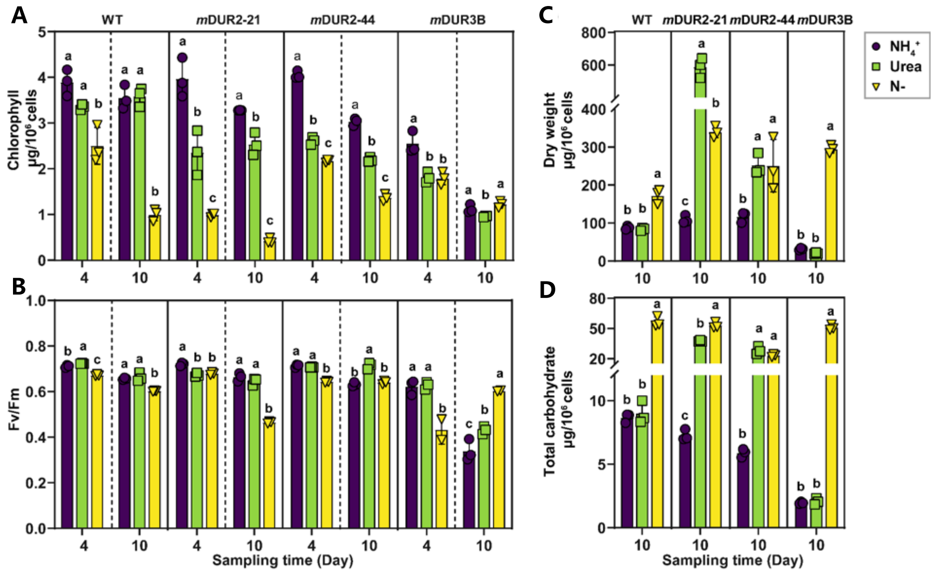


Figure S5. **The physiological response to DUR2 and DUR3B inactivation in *C. reinhardtii* grown under different nitrogen conditions. (A)** The contents of total cellular chlorophyll; **(B)** *F*v/*F*m ratio; **(C)** Dry weight; **(D)** Total cellular carbohydrate. Asterisks and different letters represent significant differences between different treatment groups ($p < 0.05$, single asterisks and letters; $p < 0.01$, doble asterisks). Note: each strain cultured in ammonium condition was used as control.

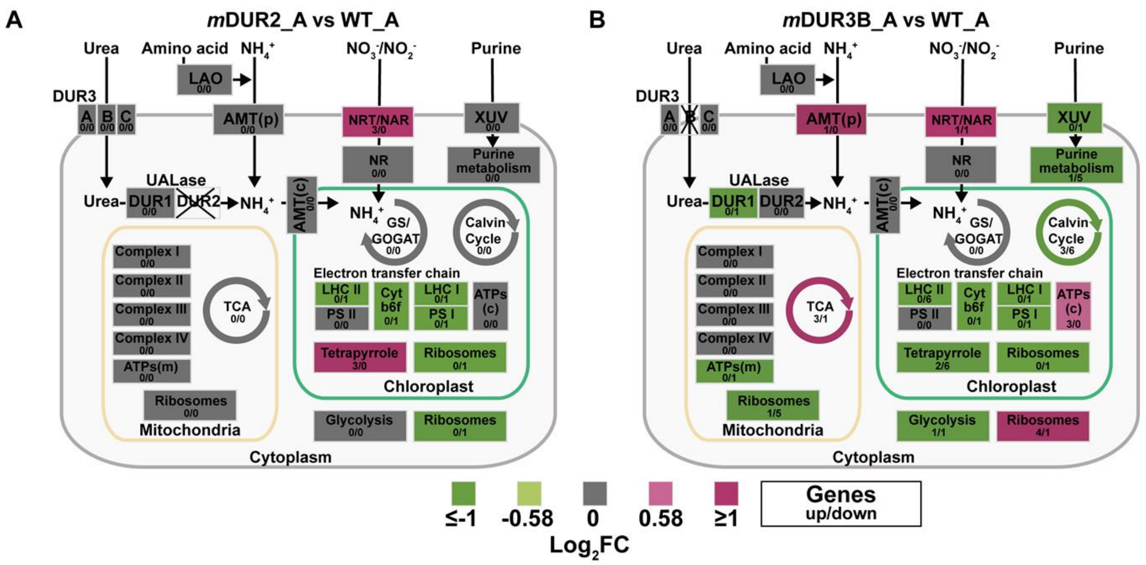


Fig. S6. **Schematic representation of metabolic pathways under different treatment conditions**. **(A)** *m*DUR2_A vs. WT_A, **(B)** *m*DUR3B_A vs. WT_A. AMT (c) and (p): ammonium transporter which located in chloroplast membrane and plasma membrane, respectively; LAO: L-amino acid oxidase; UC: urea carboxylase; AH: allophanate hydrolase; XUV: related to purine transporters; NRT/NAR: nitrate/nitrite transporter; DUR3, urea transporters; LHC I and LHC II: light-harvesting complex I and II; PS I and PS II: photosystem I and II protein; ATPs (c) and ATPs (m): chloroplast and mitochondria ATP synthase; NR: nitrate reductase; GS/GOGAT: glutamine glutamate cycle; TCA: tricarboxylic acid cycle; Tetrapyrrole: tetrapyrrole synthesis; Complex I - IV: mitochondrial respiratory chain complex I - IV. All the DGEs were listed in Table S11.

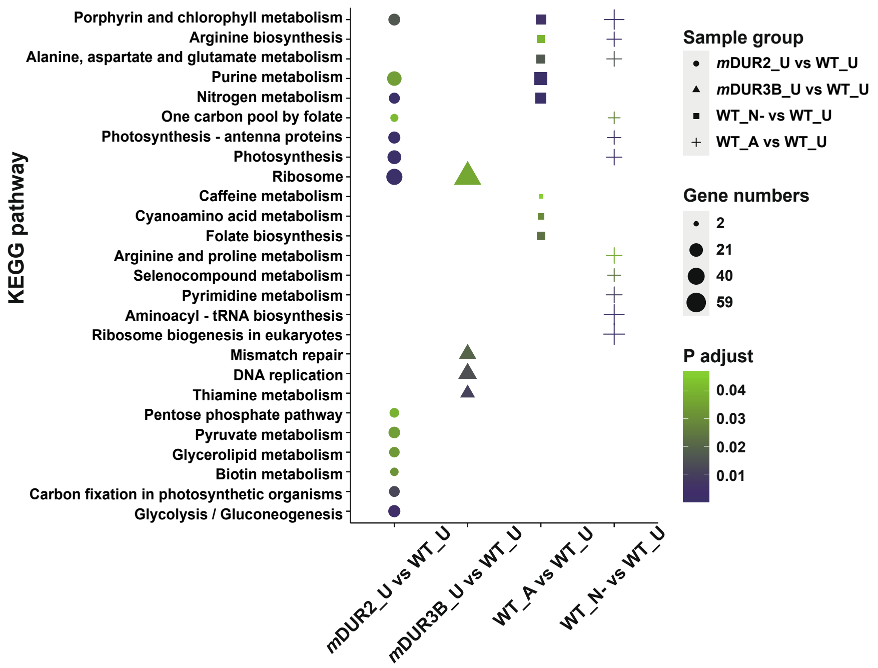


Figure S7. **Significant enriched Kyoto Encyclopedia of Genes and Genomes (KEGG) pathways.** WT_A, WT_U, and WT_N- represent wild type *C. reinhardtii* cultured in $NH_4^+$, urea, and nitrogen deficiency conditions, respectively; *m*DUR2_U and *m*DUR2_N- represent *DUR2* mutant strain cultured in urea conditions and nitrogen deficient conditions; *m*DUR3B_U and *m*DUR3B_N- represent *DUR3B* mutant strain cultured in urea and nitrogen deficient conditions.

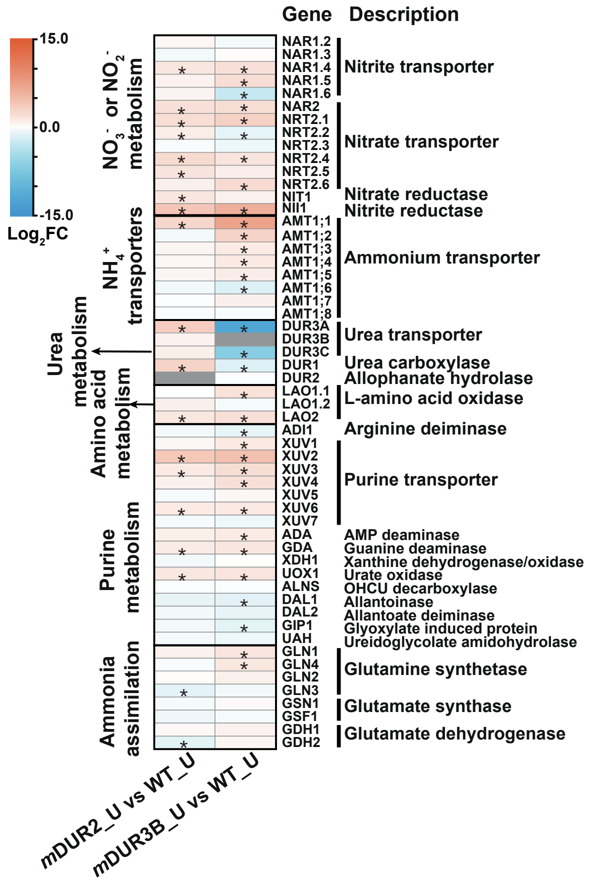


Figure S8. **Expression profiles of genes involved in nitrogen metabolism.** The red-blue heat maps were drawn according to $log_2FC$. Red and color indicates increased and decreased relative mRNA abundances, respectively. Asterisks represent significantly expressed differential genes.

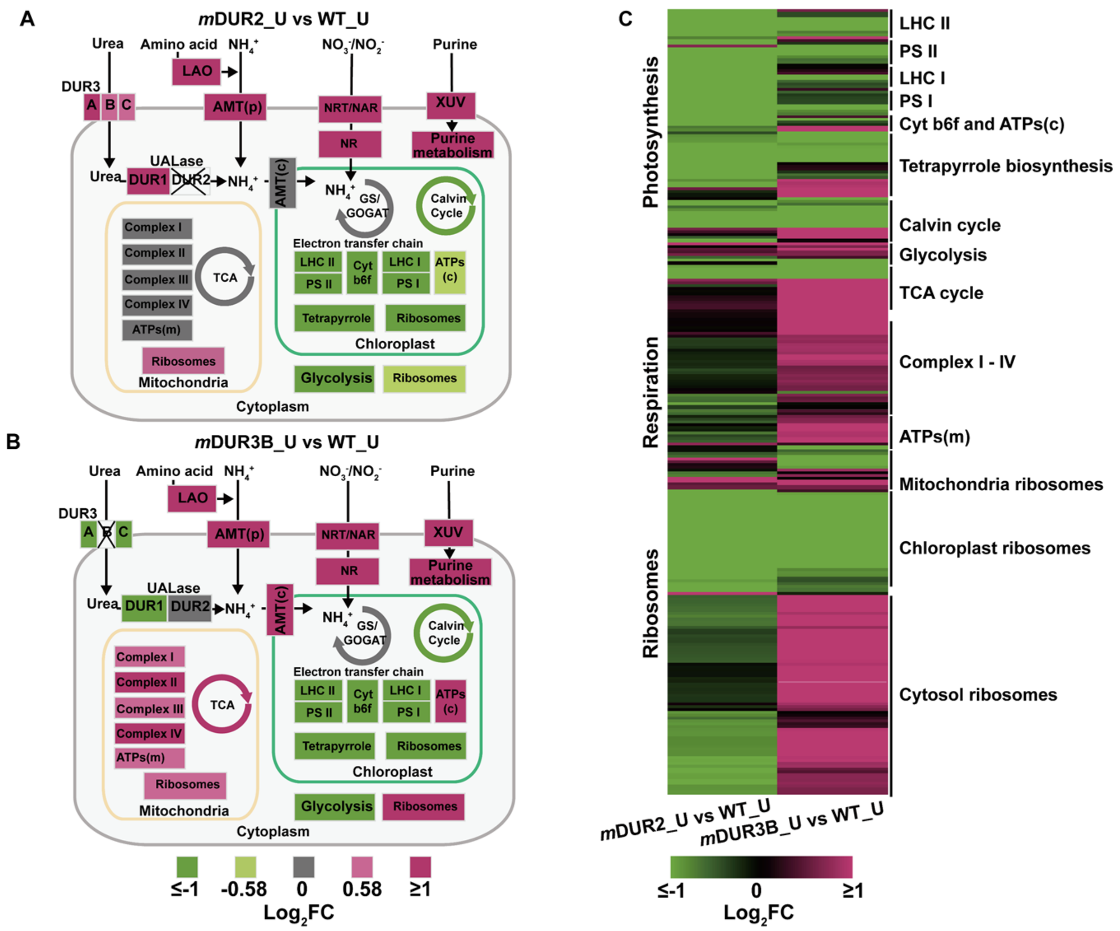


Figure S9. **Schematic representation of metabolic pathways after DUR2 and DUR3B inactivation under different nitrogen treatment groups**. **(A)** *m*DUR2_U vs. WT_U; **(B)** *m*DUR3B_U vs. WT_U; **(C)** Expression profiles of DEGs in A and B. Color bar represents the normalized gene expression level. AMT (c) and (p): ammonium transporter which located in chloroplast membrane and plasma membrane, respectively; LAO: L-amino acid oxidase; UC: urea carboxylase; AH: allophanate hydrolase; XUV: related to purine transporters; NRT/NAR: nitrate/nitrite transporter; DUR3, urea transporters; LHC I and LHC II: light-harvesting complex I and II; PS I and PS II: photosystem I and II protein; ATPs (c) and ATPs (m): chloroplast and mitochondria ATP synthase; NR: nitrate reductase; GS/GOGAT: glutamine glutamate cycle; TCA: tricarboxylic acid cycle; Tetrapyrrole: tetrapyrrole synthesis; Complex I - IV: mitochondrial respiratory chain complex I - IV. The DGEs of nitrogen metabolism was showed in Fig. 6B. All the DGEs were listed in Table S9 and S10.
